# A mechanistic hydro-epidemiological model of liver fluke risk

**DOI:** 10.1101/307348

**Authors:** Ludovica Beltrame, Toby Dunne, Hannah Rose Vineer, Josephine G. Walker, Eric R. Morgan, Peter Vickerman, Catherine M. McCann, Diana J.L. Williams, Thorsten Wagener

## Abstract

The majority of existing models for predicting disease risk in response to climate change are empirical. These models exploit correlations between historical data, rather than explicitly describing relationships between cause and response variables. Therefore, they are unsuitable for capturing impacts beyond historically observed variability and cannot be employed to assess interventions. In this study, we integrate environmental and epidemiological processes into a new mechanistic model, taking the widespread parasitic disease of fasciolosis as an example. The model simulates environmental suitability for disease transmission, explicitly linking the parasite life-cycle to key weather-water-environment conditions. First, using epidemiological data, we show that the model can reproduce observed infection levels in time and space over two case studies in the UK. Second, to overcome data limitations, we propose a calibration approach based on Monte Carlo sampling and expert opinion, which allows constraint of the model in a process-based way, including a quantification of uncertainty. Finally, comparison with information from the literature and a widely-used empirical risk index shows that the simulated disease dynamics agree with what has been traditionally observed, and that the new model gives better insight into the time-space patterns of infection, which will be valuable for decision support.

## 1. Introduction

The transmission of several highly pathogenic infectious diseases is closely linked to weather and environmental conditions (1). Waterborne diseases, such as cholera, are directly affected by hydrometeorological factors such as rainfall, through transport and dissemination of the pathogens, and water temperature, through their development and survival rates. Diseases involving a vector or intermediate host as part of their life-cycle, such as schistosomiasis, are controlled by characteristics of the water environment and land surface also indirectly, through their influence on the vector or host (2,3).

Our environment is changing at unprecedented rate due to climate change and direct human activities (4,5), with implications for the behaviour, lifespans and distribution of these diseases and their carriers (6,7). Evidence of climate and environmental-driven changes in the phenology of pathogens and incidence of diseases already exists. The increase in frequency and intensity of extreme weather events is altering the occurrence of floods and droughts, changing the concentration of infectious agents such as *Vibrio cholerae* in the water environment and human exposure to infection (3). Similarly, changes in the prevalence of schistosomiasis have been observed due to the expansion of the snail intermediate host habitat, following the construction of dams and implementation of irrigation schemes to meet demands for food and energy from increasing numbers of people (8).

As climate change accelerates and other human-caused disturbances increase, it is urgent to assess impacts on disease transmission, to guide interventions to reduce and/or mitigate risk (9). To this end, we need to: (a) understand the mechanisms by which the environment affects the epidemiological processes, addressing the system as a whole, (b) represent these processes with models that are explicit in space and time, to more reliably simulate conditions beyond historically observed variability, and (c) test these models in new ways, since simply reproducing past observations may no longer be sufficient to justify their use for decision support (1,3,7,10–12).

However, most current models for predicting changes to disease patterns in response to climate change are empirical (7,13,14). This means they do not explicitly represent mechanisms, but are based on statistical correlations between historical data, thus becoming unreliable when extrapolated to novel conditions, e.g. into different regions or future climates (15). Moreover, empirical models do not allow for what-if analyses, i.e. they cannot be used to test the effect of interventions on disease incidence, which would be valuable for supporting decision-making (10,16).

In this paper, we incorporate knowledge of environmental and epidemiological processes into a new integrated mechanistic model, using fasciolosis as an example. This is a globally distributed parasitic disease of livestock and zoonosis, whose most widespread agent is *Fasciola hepatica*, the common liver fluke (17). Clinical signs of disease in animals include weight loss, anaemia and sudden death, while sub-clinical infections result in lowered productivity and are estimated to cost the livestock industry $3 billion per year, globally (18,19). Risk of infection with liver fluke is strongly influenced by weather and environmental conditions, especially temperature and soil moisture, as the parasite has an indirect life-cycle involving an intermediate host (in the case of *F. hepatica*, the amphibious mud snail *Galba truncatula*) and free-living stages, which grow and develop in the environment (20–22).

Addressing fasciolosis is urgent for a number of reasons. First, resistance to available antiparasitic drugs is on the rise worldwide, making disease control challenging (23). Secondly, increases in disease prevalence, expansions into new areas and shifts in its seasonality have been observed in recent years and attributed to altered temperature and rainfall patterns, raising concerns about the effects of climate change in the future (23,24). Finally, it is emerging as a major disease in humans, with at least 2.4 million people infected around the world, and human treatment relying on the same veterinary drug to which resistance is increasing (25). Climate-based fluke risk models have been developed since the 1950s (20,26–28). The Ollerenshaw Index is the best-known example and is still actively used to predict disease severity in Europe (20,29,30). However, these models are empirical and therefore of little use for assessing risk under changing conditions. On the other hand, previous attempts to model fasciolosis mechanistically, in connection with climate, neglect the role of soil moisture dynamics in driving infection and do not account for the spatial aspect of the disease (e.g. 19,31).

Therefore, in this study, we introduce a new mechanistic coupled hydro-epidemiological model for liver fluke, which explicitly represents the parasite life-cycle in time and space, linked with key environmental conditions. We then parameterise the model for two case studies in the UK and assess whether it can replicate temporal and spatial variability of observed infection levels. To overcome limitations of available epidemiological data, we propose a calibration approach that combines observations and expert knowledge. Finally, we further evaluate the model by comparing it with the widely-used empirical Ollerenshaw Index.

## 2. The Hydro-Epidemiological model for Liver Fluke

The Hydro-Epidemiological model for Liver Fluke (HELF) quantitatively captures the mechanisms underlying transmission of fasciolosis, explicitly describing the causal relationships between hydrometeorological factors and biological processes, instead of simply relying on correlation. To this end, HELF integrates TOPMODEL (32,33), an existing hydrological model which we use to simulate soil moisture dynamics, and a novel epidemiological model, which represents the parasite life-cycle. TOPMODEL is chosen because its underlying assumptions are physically realistic for humid-temperate catchments, such as UK catchments, where the dominant mechanism of runoff generation is surface saturation (32). The epidemiological model is developed based on current understanding of the life-cycle of *Fasciola hepatica* and its dependence upon soil moisture and temperature (20–22).

### 2.1. Hydrological component

TOPMODEL is a catchment-scale rainfall-runoff model, which has been extensively used in humid-temperate areas for different applications (e.g. references in 34). The model uses temperature, rainfall and Digital Elevation Model (DEM) data to estimate, at each time step, spatially distributed soil moisture over the catchment (calculated as a saturation deficit), as well as streamflow at the catchment outlet. The model we use is based on the version explained in (33) and has seven parameters (Table 1).

**Table 1:**
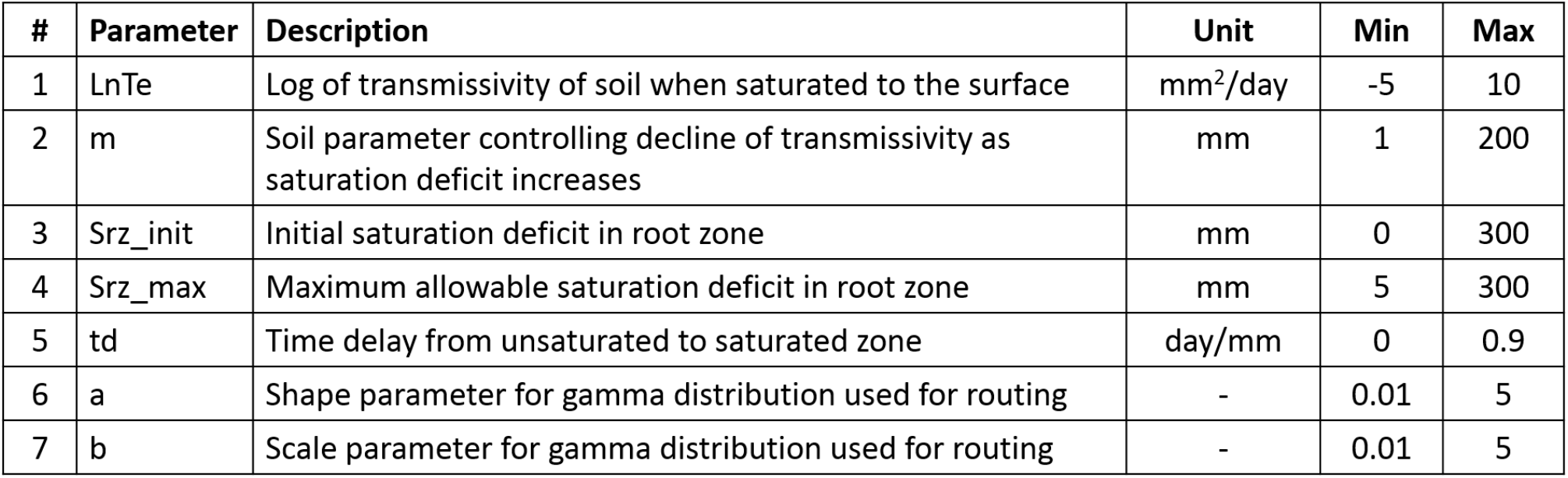
Hydrological model parameters and initial ranges.

| # | Parameter | Description | Unit | Min | Max |
| --- | --- | --- | --- | --- | --- |
| 1 | LnTe | Log of transmissivity of soil when saturated to the surface | mm<sup>2</sup>/day | -5 | 10 |
| 2 | m | Soil parameter controlling decline of transmissivity as saturation deficit increases | mm | 1 | 200 |
| 3 | Srz_init | Initial saturation deficit in root zone | mm | 0 | 300 |
| 4 | Srz_max | Maximum allowable saturation deficit in root zone | mm | 5 | 300 |
| 5 | td | Time delay from unsaturated to saturated zone | day/mm | 0 | 0.9 |
| 6 | a | Shape parameter for gamma distribution used for routing | - | 0.01 | 5 |
| 7 | b | Scale parameter for gamma distribution used for routing | - | 0.01 | 5 |

In TOPMODEL, hydrological processes through the soil are represented using a sequence of conceptual stores for which the model estimates water balances. An interception store, representing vegetation cover, must be filled by rainfall before infiltration into the soil can occur. When water infiltrates into the soil, it first enters the root zone, from which it evaporates as a function of potential evapotranspiration, maximum capacity of the store, and its actual water content. Water that is not evaporated or retained by the soil percolates to the saturated zone (i.e. the groundwater), which contributes to the channel network through subsurface flow.

To simulate the spatial distribution of soil water content over the catchment, the water balance accounting routine described, which is lumped at the catchment scale, is integrated with spatially distributed topographic information derived from DEM data. The effect of topography is captured for each grid cell through calculation of a Topographic Index: 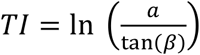, where *a* is the upslope contributing area and tan(*β*) the local slope. The index is used as a measure of the likelihood that a grid cell becomes saturated by downslope drainage: high values occur over flat areas in valleys, which tend to saturate first, whereas low values are associated with areas at the top of hills, where there is little upslope area and slopes are steep (Figure 1). The model assumes that all points with the same index value will respond similarly, hydrologically. For computational efficiency, the distribution of TI values is then discretised into classes, so that computations are performed for each class instead of for each grid cell.

**Figure 1:**
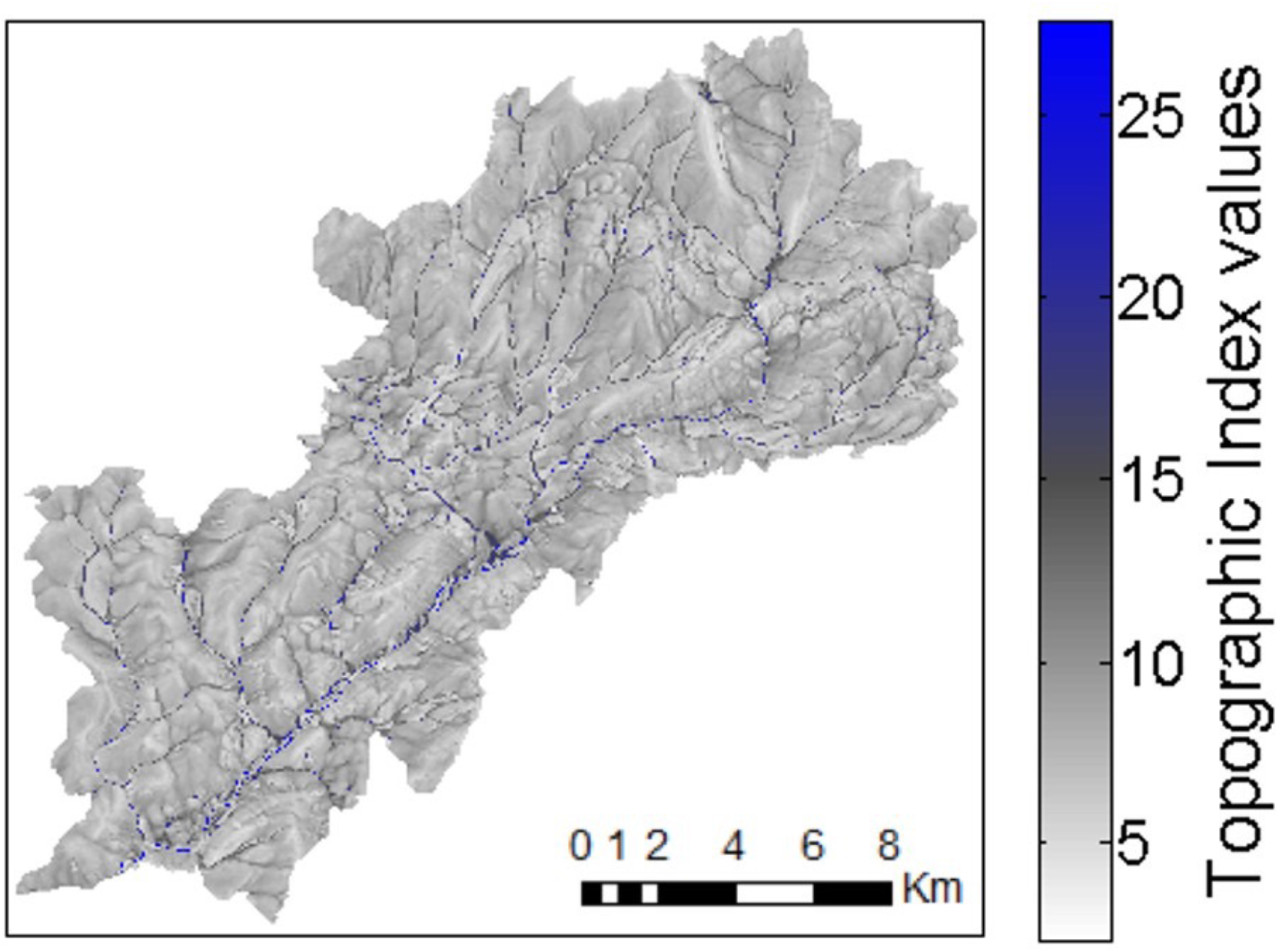
Spatial pattern of Topographic Index values for the River Tawe Catchment (UK).

Therefore, a saturation deficit for each TI class is calculated as a function of the catchment average saturation deficit, updated at each time step by water balance calculation, and the spatial distribution of the TIs. Rainfall that falls on saturated areas (i.e. where deficit is less than or equal to zero) cannot infiltrate into the soil and generates saturation-excess overland flow. Finally, total streamflow is calculated as the integrated subsurface flow and saturation-excess overland flow, and a gamma distribution is used to model the time delay in discharge generation at the catchment outlet, due to water moving through the river channel.

### 2.2 Epidemiological component

The epidemiological model component of HELF represents the stages of the liver fluke life-cycle that live on pasture: eggs, miracidia, snail infections and metacercariae (Figure 2). Development and survival of these, as well as the presence of mud snails, require particular temperature conditions and wet soil. Therefore, the model takes as input variables temperature and soil moisture, as well as an egg scenario (i.e. number of embryonic eggs we assume are deposited on each TI class at each time step by infected animals), to calculate the abundance of individuals in each life-cycle stage.

**Figure 2:**
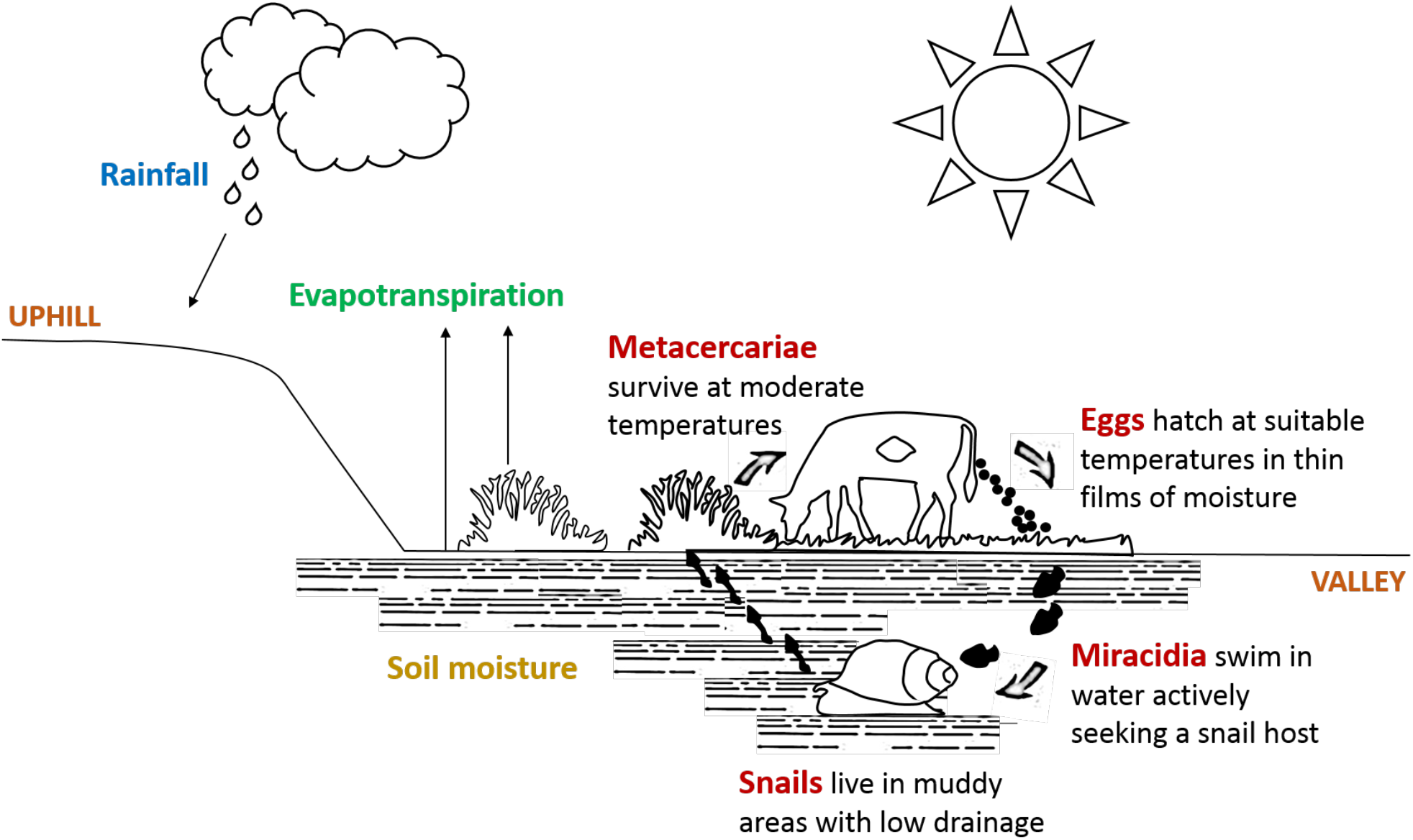
Simplified representation of the liver fluke life-cycle, with an amphibious mud snail serving as intermediate host.

Once passed out on pasture in the faeces of infected animals, Eggs (E) develop at a temperature-dependent rate, and hatch into miracidia when both temperature and soil moisture conditions are suitable (35). Miracidia (Mi) are short lived: either they find a snail host or die within 24 hours (35,36). Therefore, progression from miracidium to the next stage is calculated as the probability of finding a snail. This is assumed to depend on soil moisture and temperature, as *Galba truncatula* snails are only found in poorly drained areas and are known to hibernate with cold weather and aestivate during hot dry periods (35). Snail infections (SI) also develop in the model as a function of both temperature and soil moisture, as development within the snail may be halted due to hibernation and aestivation (21), until parasites emerge from snails in the form of cercariae. Once attached to grass as Metacercariae (Me), these survive on pasture and retain infectivity based on temperature, with moderate weather being most favourable (35).

Each stage, except for miracidia that only have a lifespan of one day, is represented as a pool of developing cohorts of individuals to capture maturation progress in a realistic way. Individuals in different cohorts are exposed to different environmental conditions, and therefore will develop at different times (35,36). We account for this by using two state variables for each cohort within each stage: number of individuals in the cohort and “age” of the cohort. The rationale is that each cohort has a certain age, which increases with the number of days that have suitable environmental conditions, until the cohort eventually matures to the next life-cycle stage. Output from a stage is then the sum of cohorts per unit area which mature to the next one.

At each time step, development and/or survival rates for a stage are calculated based on the value of the relevant environmental conditions for that stage at that time step, and on the stage-specific requirements for development/survival, which are model parameters (Table 2). The technique employed to build the functions to calculate these rates has previously been used for modelling both liver fluke and other parasites (e.g. 19,31). For temperature-dependent rates, we use information in the literature from laboratory experiments or controlled micro-environment studies that examine the time to development or death at a range of constant temperatures. First, rates are calculated for each constant temperature from the reported e.g. time to development (i.e. rate = 1/time to development); then piecewise linear models are fitted to these rates, yielding a regression equation which can be used to estimate the daily rates based on the time series of observed temperature. For soil moisture, we adopt the same approach, assuming that development is fastest when the soil is saturated to the surface (i.e. when deficit = 0) and that there is no development above a certain maximum deficit (20,35). For stages with both temperature and soil moisture requirements, we allow for development to progress as a function of both (Figure 3).

**Table 2:** Epidemiological model parameters and initial ranges.

| # | Parameter | Description | Unit | Min | Max | References |
| --- | --- | --- | --- | --- | --- | --- |
| 1 | S1MinTemp | Min temp for egg development | °C | 5 | 15 | 20, 21, 35 |
| 2 | S1PeakTemp | Optimal temp for egg development | °C | 15 | 27 | 35, 36 |
| 3 | S1PeakRate | Egg development rate at optimal temp | day<sup>-1</sup> | 0.025 | 0.5 | 35, 36 |
| 4 | S1MortRate | Egg mortality rate | day<sup>-1</sup> | 0 | 0.0693 | 35 |
| 5 | S1MaxTemp | Max temp for egg development | °C | 27 | 40 | 35 |
| 6 | S1TrigSatDef | Max saturation deficit for egg hatching | mm | 1 | 100 | 21, 35 |
| 7 | S1TrigMinTemp | Min temp for egg hatching | °C | 5 | 15 | 21 |
| 8 | S2SatDef | Max saturation deficit for miracidia finding snails | mm | 1 | 100 | 20, 35, 36 |
| 9 | S2SatSlope | Slope defining optimal saturation deficit for miracidia finding snails | - | 0.01 | 0.5 |  |
| 10 | S2MinTemp | Min temp for miracidia finding snails | °C | 5 | 15 | 35 |
| 11 | S2PeakTemp | Optimal temp for miracidia finding snails | °C | 15 | 27 | 35 |
| 12 | S2MaxTemp | Max temp for miracidia finding snails | °C | 27 | 40 | 21 |
| 13 | S3SatDef | Max saturation deficit for snail infections development | mm | 1 | 100 | 20, 35, 36 |
| 14 | S3SatSlope | Slope defining optimal saturation deficit for snail infections development | - | 0.01 | 0.5 |  |
| 15 | S3MinTemp | Min temp for snail infections development | °C | 5 | 15 | 20, 21, 30 |
| 16 | S3PeakRate | Snail infections development rate at optimal temp | day<sup>-1</sup> | 0.0204 | 0.0357 | 35 |
| 17 | S3MortRate | Snail infections mortality rate | day<sup>-1</sup> | 0 | 0.0693 | - |
| 18 | S4MinTemp | Min temp for metacercariae survival | °C | -20 | 10 | 35 and therein |
| 19 | S4PeakTemp | Optimal temp for metacercariae survival | °C | 10 | 15 | 35 and therein |
| 20 | S4MaxTemp | Max temp for metacercariae survival | °C | 15 | 40 | 35 and therein |
| 21 | S4PeakRate | Metacercariae survival rate at optimal temp | day<sup>-1</sup> | 0.0027 | 0.0333 | 35 and therein |
| 22 | S4MinRate | Metacercariae survival rate at min/max temp | day<sup>-1</sup> | 0.0333 | 0.5 | 35 and therein |

**Figure 3:**
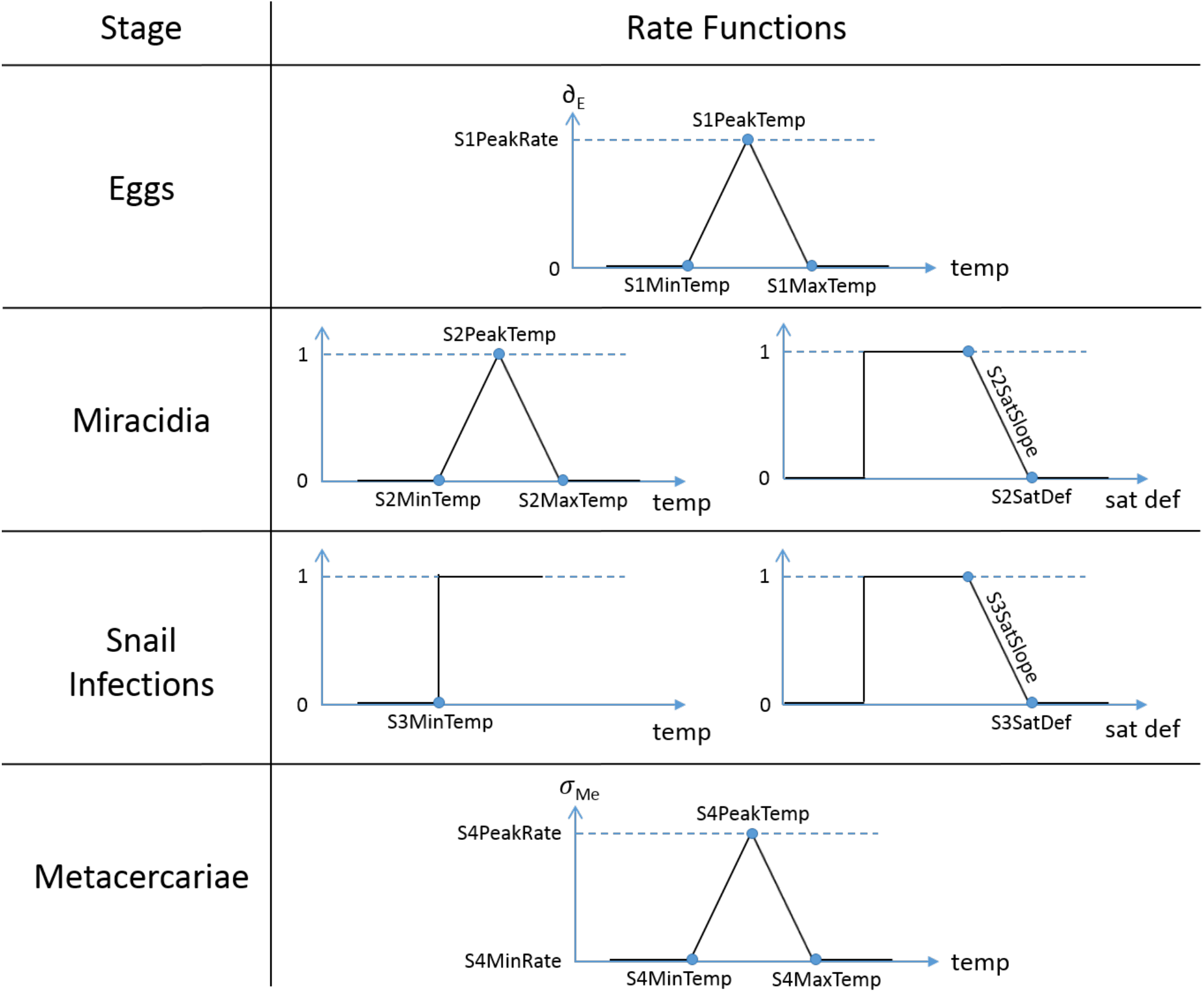
Functions used in HELF to calculate temperature and soil moisture-dependent development and mortality rates.

### 2.3. Coupled model

The coupled hydro-epidemiological model runs with a daily time step and has a total of 29 parameters. For each day, HELF calculates the catchment average saturation deficit based on rainfall and temperature, and derives the saturation deficit for each of 25 TI classes, based on this and the TI value for the class. Then, for each class and life-cycle stage, the model calculates the relevant development and/or survival rates, based on environmental conditions. The age of each cohort is updated based on the development rate, and, given an egg scenario, the model finally computes the number of individuals in the stage as a function of the number from the previous time step, plus the sum of the cohorts developed from the previous stage, minus those that die (Figure 4). Therefore, the model outputs are the abundances of developed eggs, snails located and infected by miracidia, developed snail infections, and infective metacercariae surviving on pasture, which represents the environmental suitability for disease transmission to grazing livestock. These variables, calculated for each TI class, are then mapped back onto each grid cell in the catchment.

**Figure 4:**
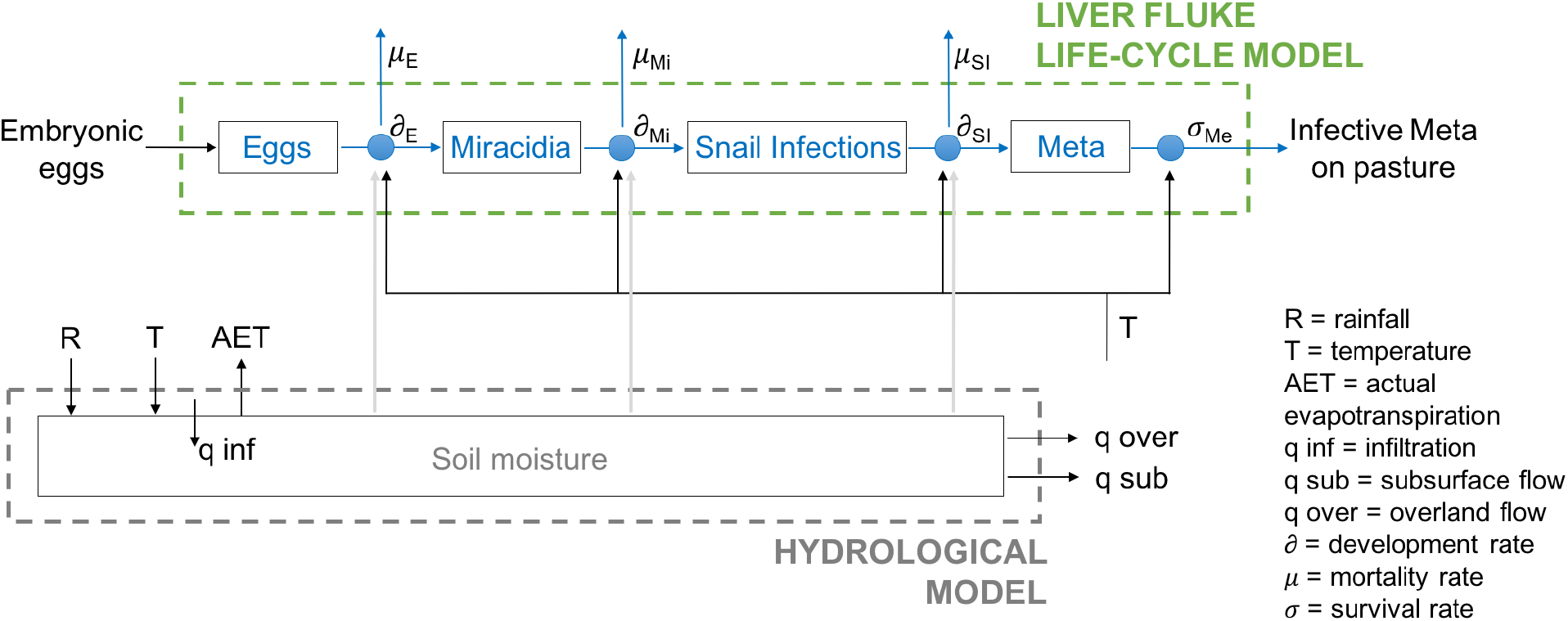
Simplified flow diagram of HELF, which integrates a hydrological and a liver fluke life-cycle component, to simulate the abundance of infective metacercariae (Meta) on pasture.

With regard to the egg scenario, the current model assumes that 100 embryonic eggs are introduced on each TI class daily, over the whole simulation period. This means we are considering a scenario of continuous livestock grazing and no disease management over the catchment. However, this assumption can be easily changed: the fact that the egg scenario is a model input leaves the end-user of the model the possibility to estimate how the environmental suitability for disease transmission translates into risk of infection based on local farm management factors such as grazing season length or disease control strategy.

## 3. Study sites and data

To test HELF, we apply it to two catchments in the UK, located in South Wales and the North-West Midlands (England), respectively. The datasets employed include hydro-meteorological and epidemiological data.

### 3.1 The Tawe and Severn Catchments

The River Tawe flows approximately 50km south-westwards from its source in the Brecon Beacons to the Bristol Channel at Swansea. The catchment is about 240km^2^ in area, with elevation ranging from about 10 to 800m a.s.l., and most of the area characterised by a relatively impermeable bedrock. The River Severn rises in mid Wales and flows through Shropshire, Worcestershire and Gloucestershire, before also discharging into the Bristol Channel. The catchment, closed at Upton on Severn, is about 6850km^2^ in area, with elevation range and geological characteristics similar to the Tawe (37). Both catchments have grassland as the dominant land use (Figure 5), which is extensively used for livestock farming, and are located in known fluke endemic areas (38). Moreover, these areas are predicted to become increasingly warmer and wetter on average (39), which suggests they will become even more favourable for liver fluke transmission in the future.

**Figure 5:**
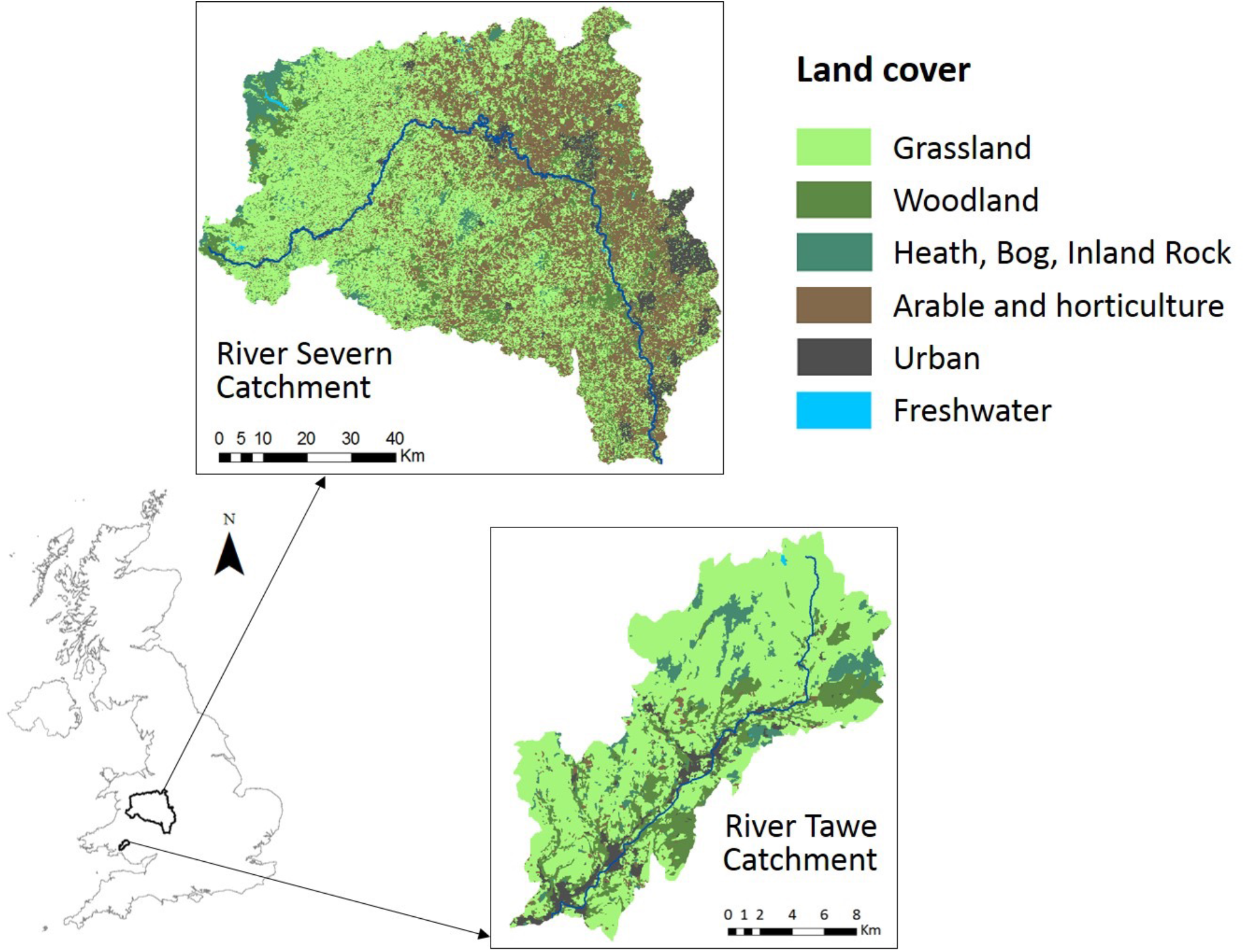
Location and land cover map of the Tawe and Severn Catchments (36).

### 3.2 Hydro-meteorological and epidemiological data

The hydro-meteorological dataset includes daily observations of rainfall, temperature and discharge. Gridded time series of rainfall and temperature are obtained from CEH-GEAR and the UK MetOffice, respectively. For both case studies, to run HELF, we take the average over the grid cells within the catchment. For the Tawe we use these time series for a 12-year period (1999-2010), whereas for the Severn we use 2 years of data (2013-2014), in line with the available epidemiological data periods. Observed discharge over the same years, at Ynystanglws for the Tawe and Upton-on-Severn for the Severn, are obtained from (37). DEM data for both catchments is obtained from NextMap with spatial resolution of 5m, then aggregated to 25m.

The epidemiological dataset consists of a time series from the Veterinary Investigation Diagnostic Analysis (VIDA) database for the Tawe and a spatial dataset based on Faecal Egg Counts (FECs) for the Severn. The VIDA database, compiled from reports from the UK Government’s Animal and Plant Health Agency regional labs, provides diagnoses of fasciolosis made from ill or dead animals. The time series we use is the monthly number of sheep diagnosed with acute fasciolosis from the post code district areas within the Tawe catchment over 1999-2010. This data is believed to reflect well the temporal dynamics of within-year infection levels, but may not always reflect the magnitude of infection in the field, as the rate of submission of animals to the labs is influenced by multiple factors (40). In our series, no cases are reported for 2001 and values over the following years are low, which may have been affected by the 2001 outbreak of foot-and-mouth disease in the UK. On the other hand, the spatial dataset for the Severn catchment consists of 174 cattle herds, from farms within a 60km x 75km area in Shropshire, that have been classified into infected and non-infected based on FECs collected over 0ct2014-Apr2015. Unlike VIDA, this is active surveillance data, and thus more likely to reflect true levels of infection. However, rather than a continuous/quantitative measure of the magnitude of infection, this dataset only provides a binary classification into positive-negative farms, at one moment in time and at a limited number of points within the catchment.

## 4. Model calibration and testing

HELF comprises parameters related to aspects of the local environment and parameters related to the phenology of the parasite (Tables 1–2). Usually, ranges of values can be derived from the literature for these, rather than point estimates, partly because of their associated natural variability and partly due to uncertainty and poor understanding. This may result in different parameter sets providing equally good representations of system behaviour, with implications in terms of predictive uncertainty and limitations to the applicability of the model (15,34). Therefore, it is crucial to constrain and evaluate HELF, if we want to use it to assess disease risk in the future.

Usually models are calibrated and validated using historic records, assuming that the data available reflect the underlying system, and that conditions in the period considered are similar to those under which the model will be used. However, this may not be sufficient if data are disinformative in some respects and/or if the purpose of the model is to simulate conditions that are significantly different to those previously observed (41).

Our strategy involves multiple datasets and methods. On one hand, we have high quality continuous data for both the meteorology and hydrology. Therefore, we calibrate and validate the hydrological component of HELF by adopting a standard split-sampling approach (41). On the other hand, given the epidemiological data limitations mentioned in Section 3.2., our approach for constraining the epidemiological model component not only uses past observations, but also expert-driven rules.

### 4.1. Calibration and testing of the hydrological component

To estimate TOPMODEL parameter values and evaluate its prediction capabilities, we perform a split-sample test using streamflow observations (years 2000-2006 for calibration and 2007-2010 for validation, with 1999 as warm-up period) (41). The Shuffled Complex Evolution (SCE-UA) optimisation method is employed to find the parameter set which maximises the coefficient of determination (R^2^) between simulations and observations on our catchments (42). The algorithm samples an initial population of parameter sets from *a priori* defined ranges (Table 1) and then evolves this population of sets to find the best performing one with respect to R^2^.

### 4.2. Calibration and testing of the epidemiological component

Using the best performing parameterization obtained for TOPMODEL, first, we fit the fluke component of HELF to the two epidemiological datasets and assess whether we can reproduce the observed patterns of infection, ignoring the data limitations discussed. Secondly, under the assumption that these data may be disinformative, and given that we ultimately want to use HELF to simulate fluke risk under changing conditions, we propose an alternative calibration approach based on Monte Carlo sampling and expert knowledge. Finally, we evaluate the model by comparing it to observations from previous studies and the commonly-used Ollerenshaw Index.

#### 4.2.1. Single-objective approach using epidemiological data

To estimate parameters of the epidemiological model for the Tawe Catchment, we fit HELF to the VIDA time series by using SCE-UA to maximise the Pearson coefficient of correlation (*r*) between simulated abundance of infective metacercariae and observed number of sheep infections. As the VIDA dataset only provides a single time series for the Tawe, we aggregate the simulated abundance of metacercariae over the catchment, taking the average across TI classes. Moreover, to account for the delay between the variable we simulate and the observations, a lag parameter is included in the optimisation process, which is allowed to vary between 0 (no delay) and +5 months (18).

Similarly, to estimate parameters for the Severn Catchment, we fit HELF to the FEC-based spatial dataset. First, we divide the area over which we have observations into sub-areas with a minimum of 15 data points each. Second, we use SCE-UA to find the parameter set which maximises *r* between the simulated percentage of grid cells at risk of infection and the observed percentage of herds infected, over each sub-area. To this end, for each parameter set, firstly, we aggregate the simulated abundance of metacercariae over months Jul-Dec2014, assuming that pasture contamination over this period will be responsible for the observed infection levels (38). Secondly, we classify the simulated abundance of metacercariae in each grid cell into two classes (no-risk and risk) by setting a threshold based on the overall observed percentage of infection.

#### 4.2.2. Monte Carlo sampling-based approach using expert opinion

Given the limitations of our epidemiological datasets, we believe that simply fitting these may not be sufficient to guarantee reliability of our new model. Moreover, if HELF is to be used to assess future disease risk, its credibility should be assessed via more in-depth evaluation of the consistency with the real-world system, instead from just comparison against historical data (11). To this end, we collect information from the literature (e.g. 24,27,30) and our perceptions, to characterise the seasonality of the liver fluke life-cycle stages in the UK over years 2000-2010. This includes shifts in seasonality experienced over this period, compared to what has been traditionally observed, driven by changing temperature and rainfall patterns, but could be adjusted to account for further changes and shifts going forwards. Then, we formalise this knowledge into a set of rules:

- Rule1: retain the parameter sets that every year predict the first month of snail presence (i.e. with positive number of snails) to happen earlier than average, if temperature is above average in January-March
- Rule2: retain the sets that every year produce higher mean number of snail infections over summer (June-August), if number of rain days over May-July is above average
- Rule3: retain the sets that every year produce higher mean number of metacercariae over autumn (August-October), if rainfall is above average and the number of days>20°C is below average over May-August
- Rule4: retain the sets that every year produce higher mean number of metacercariae over winter (January-February), if total number of days>10°C is above average over Jan-Feb

Finally, we sample 8000 parameter sets using uniform distributions from ranges in Table 2, and reject all the sets producing model outputs that are inconsistent with these rules.

#### 4.2.3. Comparison with the Ollerenshaw Index

To further evaluate HELF, we use the behavioural parameterisations, i.e. those retained from sequential application of the rules, and compare disease risk simulated using these with the Ollerenshaw Index. This, calculated at the monthly scale based on number of rain days, rainfall and temperature as in (29), is the current standard for providing liver fluke forecasts in the UK, where it is used by the National Animal Disease Information Service to warn farmers about high risk years (30).

## 5. Results

### 5.1. Performance of the hydrological model

Comparison of simulated and observed daily streamflow shows that TOPMODEL is able to reproduce well the temporal dynamics of observations, including the peaks and recession periods of the hydrograph, with R^2^=0.87 during calibration and 0.84 in validation (Figure 6).

**Figure 6:**
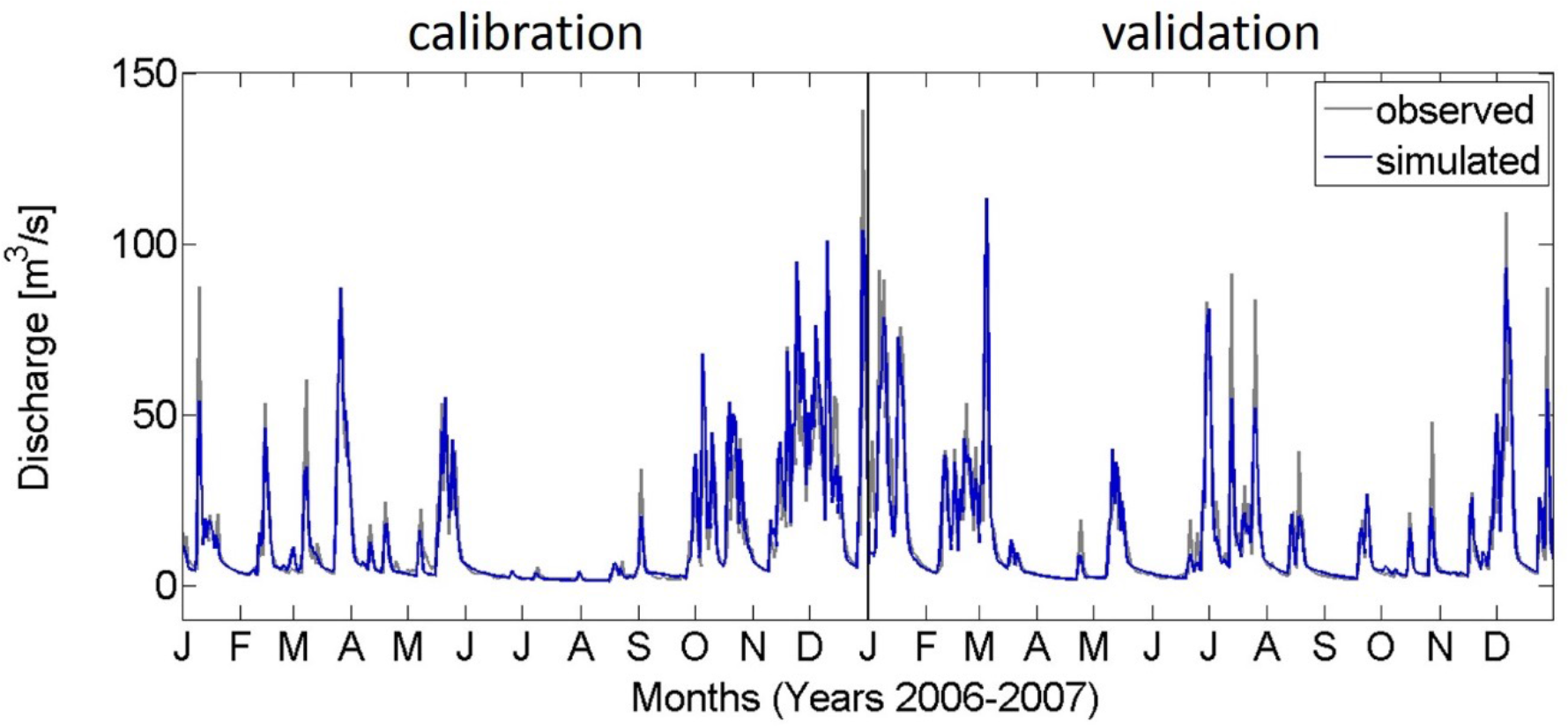
Extract of the calibration and validation periods using daily streamflow data for the River Tawe Catchment (total period is 2000-2006 for calibration and 2007-2010 for validation).

### 5.2. Performance of the epidemiological model

#### 5.2.1. Fit to epidemiological data

A delay is evident between simulated catchment average number of metacercariae and reported number of sheep diagnosed with fasciolosis from the Tawe Catchment (Figure 7). This is due to the time-lag between pasture contamination, which HELF simulates, and infection diagnosed in the animal, which the VIDA dataset reports. Except for the year 2000, for which the model predicts risk of infection that is not reflected in the VIDA numbers over 2001, HELF seems to adequately predict the observed temporal dynamics of infection. It simulates low pasture contamination for most of the period and captures the higher peaks over winters 2008-2009 and 2009-2010, driven by the preceding exceptionally wet summers and rainy autumns. The highest correlation between the two series (*r*=0.62) is found at a lag of three months, which corresponds to the prepatent period of fasciolosis reported in the literature (18). If, instead of using the whole dataset for calibration, we perform a 5-fold cross-validation, mean correlation results 0.52 in calibration and 0.41 in validation.

**Figure 7:**
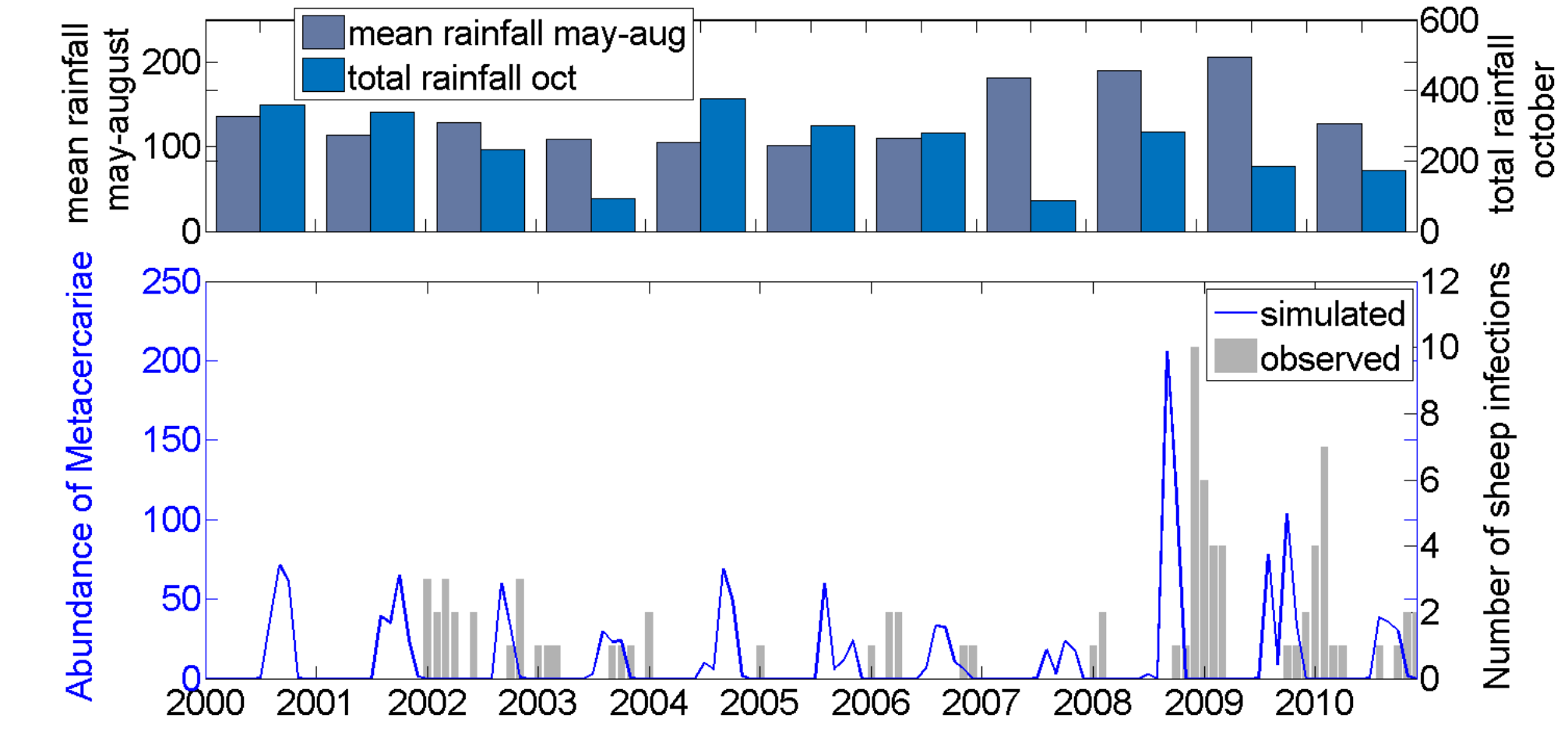
(bottom) Monthly comparison of simulated catchment average number of metacercariae and observed number of infections (VIDA data) over years 2000-2010 for the Tawe Catchment. (top) Years 2008 and 2009 have the highest mean summer rainfall within the simulation period, as well as a sufficiently wet autumn, resulting in high suitability for disease transmission.

Division of the area for which we have observations within the Severn Catchment into sub-areas with at least 15 data points each, results into 9 sub-areas (Figure 8). If we then compare the simulated percentage of grid cells at risk of infection with the observed percentage of infected herds, in each of the sub-areas, the two are in good agreement (*r*=0.83), suggesting that the model can replicate the observed spatial pattern (here, performing a leave-one-out cross-validation results in a mean absolute error of 0.1). Risk of infection seems overestimated in sub-areas A2 and A5. However, these were significantly drier than the other sub-areas in 2014 (Figure S1), and have a lower percentage of area suitable for snail hosts in terms of soil pH (Figure S2), which HELF currently does not account for.

**Figure 8:**
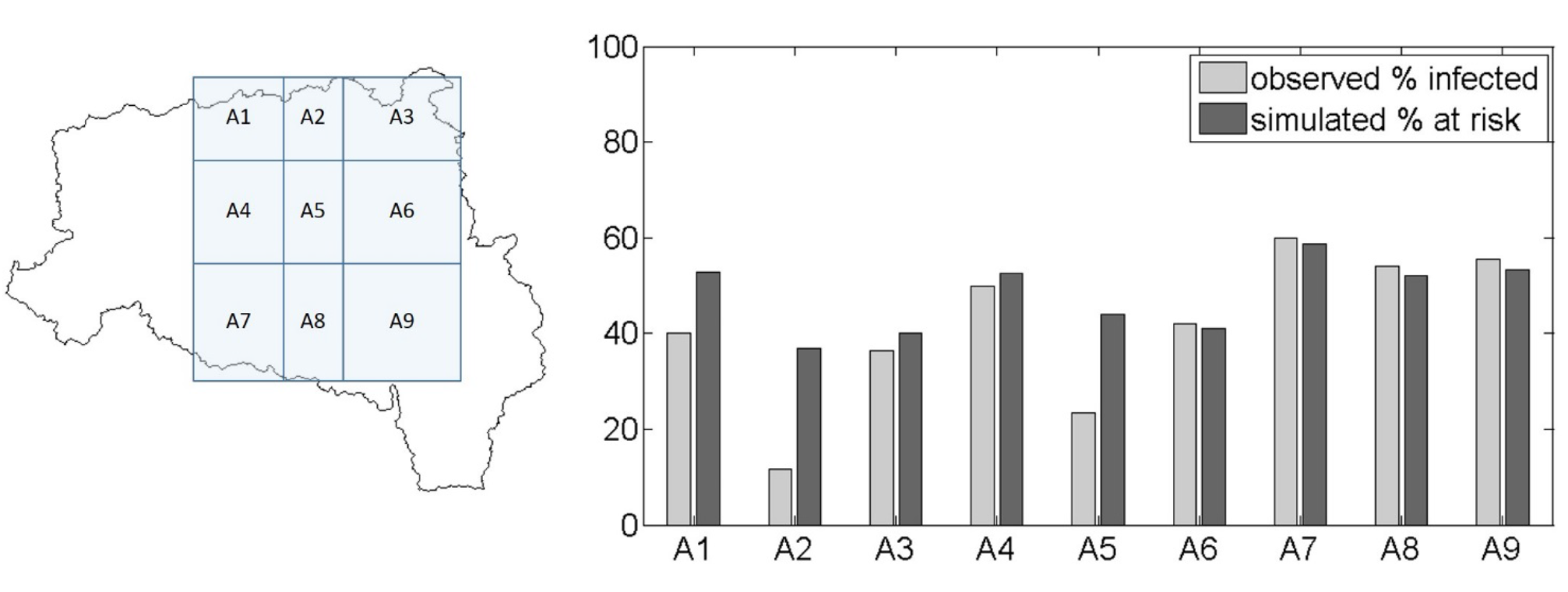
(left) Sub-areas within the Severn Catchment for which we have data points (i.e. cattle herds classified into infected and not-infected based on FECs collected over winter 2014-2015). (right) Comparison of simulated percentage of grid cells at risk of infection and observed percentage of infected herds for each sub-area.

#### 5.2.2. Result of the expert-driven approach

Sequential application of the expert-driven rules reduces the initial sample of 8000 parameter sets to 14 behavioural parameterisations (Figure 9). The resulting simulated life-cycle stages show that the abundance of developed eggs on pasture tends to increase in March, as the weather warms up, before decreasing gradually over the following months, as hatching into miracidia begins (Figure 10). Snail activity, and therefore infection of snails by miracidia, also seems to start in spring, and carries on until November, when frosts may send snails back into hibernation; whereas development of intra-molluscan infections peaks around August, leading to high numbers of infective metacercariae surviving on pasture in Autumn. Finally, if we compare the abundance of metacercariae – obtained using the whole set of behavioural parameterisations – with the VIDA time series, first, we still see the expected delay between simulations and observations (Figure 11). Moreover, we note that, on one hand, uncertainty is large in terms of magnitude of the yearly peak of infection. On the other hand, uncertainty bounds are narrower in terms of timing and duration of the outbreaks, with the number of infective metacercariae on pasture beginning to increase in July, reaching a peak in September, before decreasing again in December, on average.

**Figure 9:**
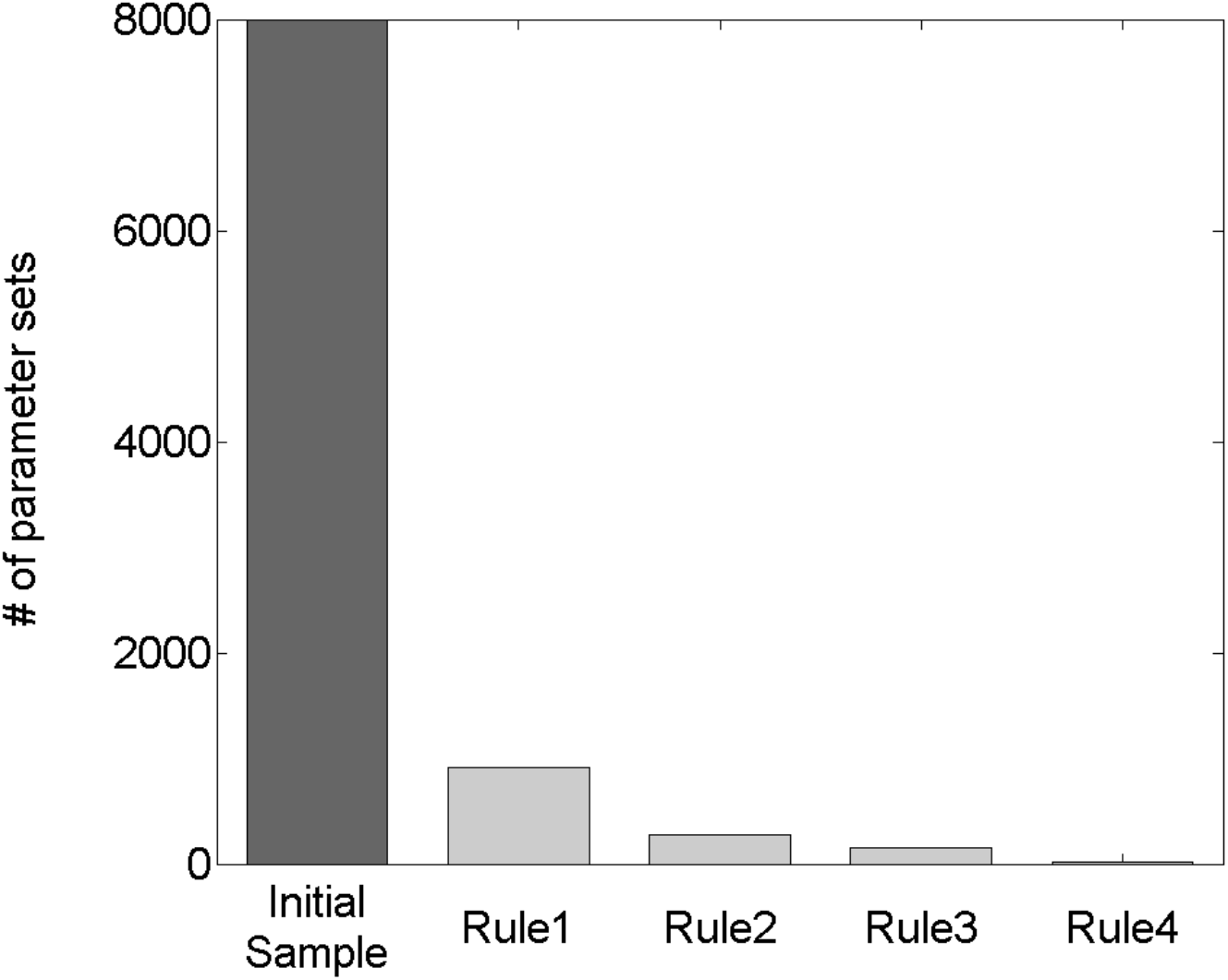
Evolution of the initial sample of 8000 parameterisations (each including the 22 epidemiological model parameters sampled from within their initial ranges) along the 4 confinement steps.

**Figure 10:**
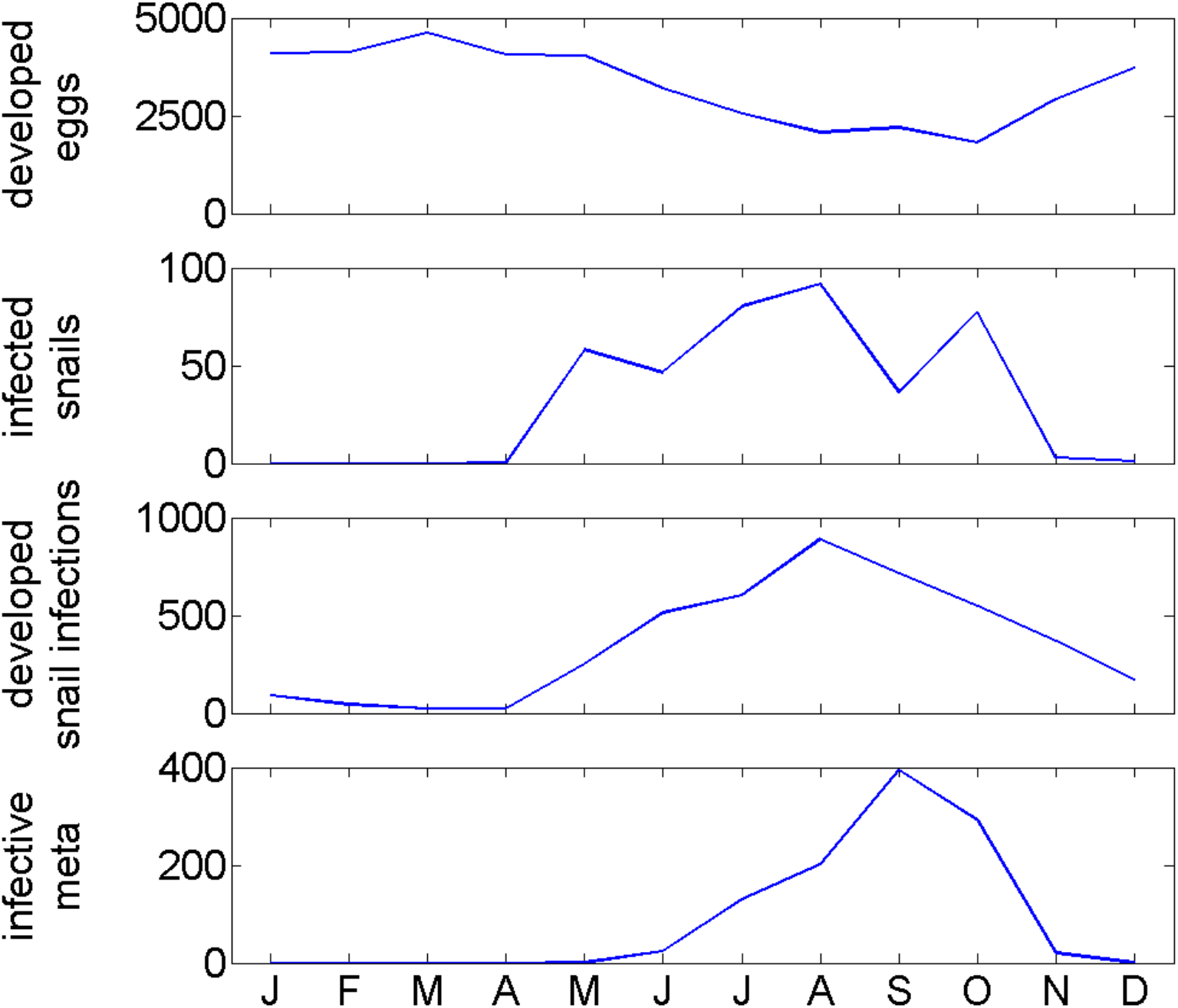
Monthly behaviour of the parasite life-cycle stages simulated with HELF for year 2001, as an example (median of the behavioural simulations).

**Figure 11:**
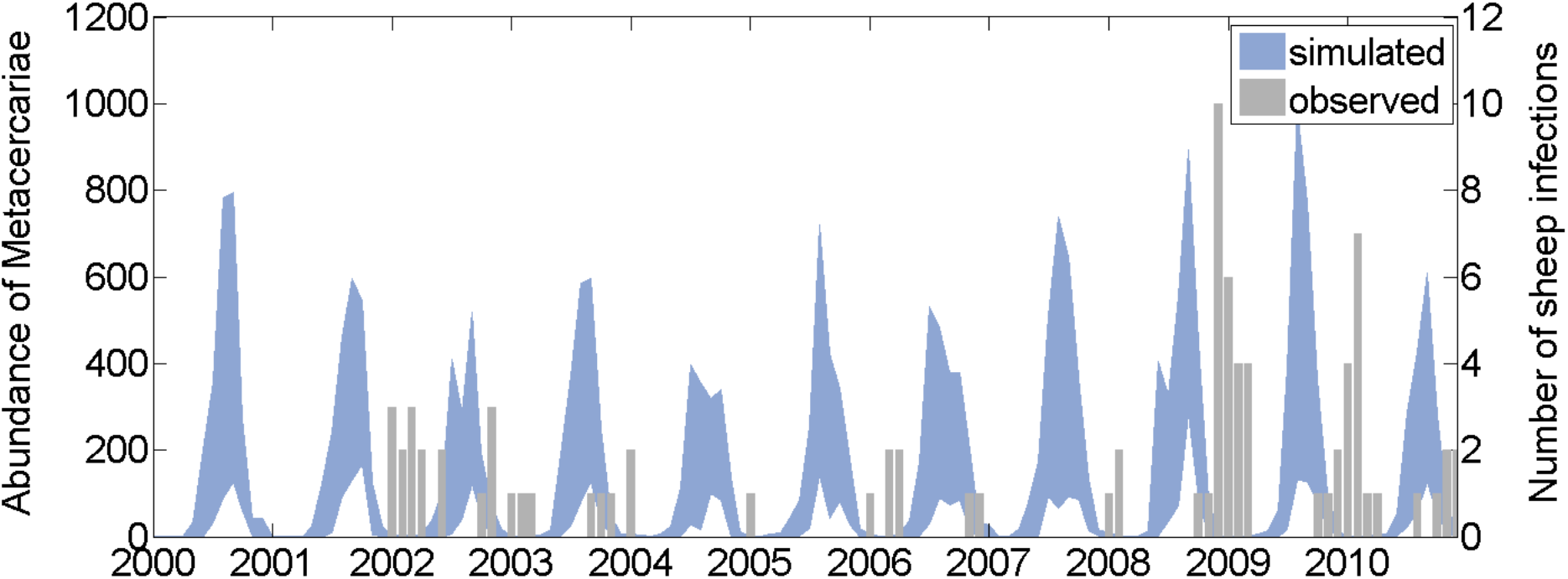
Monthly comparison of simulated catchment average number of metacercariae, obtained using the behavioural parameter sets (90% bounds), and observed number of infections (VIDA data) over years 2000-2010 for the Tawe Catchment.

#### 5.2.3. Comparison with the Ollerenshaw Index

Temporal comparison of the suitability for disease transmission simulated by HELF, constrained using the rules, with the Ollerenshaw Index, shows a time-lag of one month between the two series (Figures 12a and S3). This is due to the two models representing different things: a risk index based on monthly temperature and rainfall characteristics in the case of Ollerenshaw, and the abundance of metacercariae, based on soil moisture and accounting for the delays in the parasite life-cycle, in the case of HELF. From here we also see that, while matching the empirical index on inter-annual variation (at lag of one month, *r*=0.73), the two models’ responses may differ at higher temporal resolution. For example, the Ollerenshaw Index reaches the same peak value in years 2007 and 2008, but risk of infection in 2007 seems lower than the following year according to HELF. Comparison of the two models in space, presented in Figure 12b for August 2006 as an example, shows the presence of high risk areas in the Tawe Catchment according to both models. However, on one hand, according to the Ollerenshaw Index, no proportion of the catchment is risk-free, and risk of infection is higher in the North-East, where rainfall levels are higher (37). In contrast, for the same month, assuming an area is at risk if its number of metacercariae is positive, HELF estimates that 17.3% of the catchment is risk-free, and that there are 134 patches at risk, spread throughout the catchment, with mean size of 1.6km^2^.

**Figure 12:**
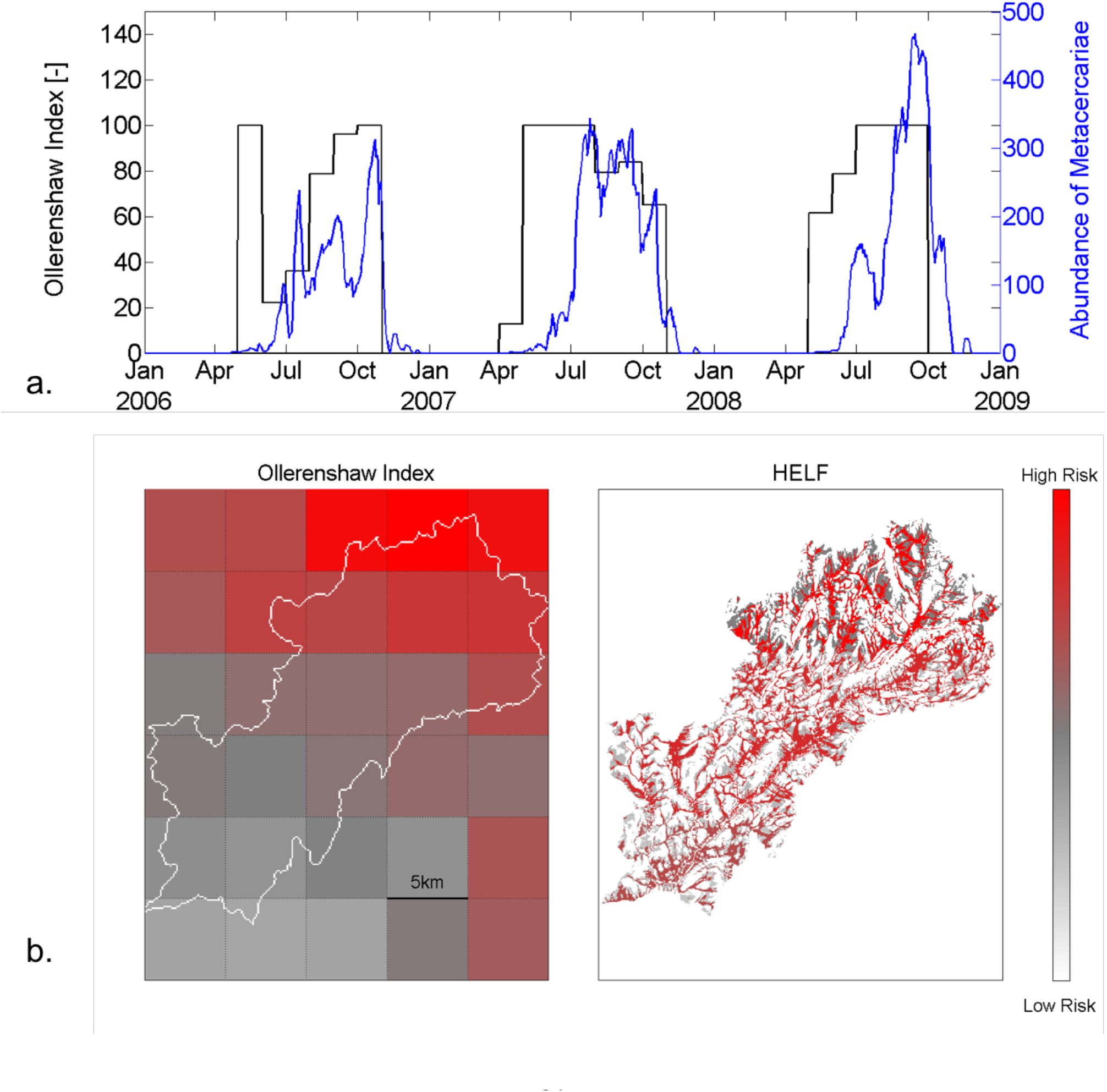
(a) Comparison of the Ollerenshaw risk index pattern, in black, with the temporal dynamics of pasture contamination simulated with HELF (median of the behavioural sets) in blue, for an extract of the simulation period over the Tawe Catchment. (b) Risk maps for August 2006, as an example, obtained using the Ollerenshaw Index (left) and HELF (right).

## 6. Discussion

In this study, we developed the first mechanistic model which simulates risk of infection with *F. hepatica* in time and space, driven by temperature and soil moisture dynamics. The novelty of our work lies in the description of the bio-physical processes underlying transmission of fasciolosis, advancing the study of the disease beyond empirical associations of infection levels with temperature and rainfall. Despite existing forecasting models calculating fluke risk based on these (20,29,30), soil moisture has always been recognised as the critical driver of disease transmission for its role on development of the free-living stages and presence of the snail intermediate hosts (20). Here we represented it using an existing hydrologic model, which is dynamic and based on spatially distributed topographic information, also known as an important fluke risk factor (27). Moreover, collaboration across the physical and biological sciences was necessary to analyse the effect of both soil moisture and temperature on the multiple parasite life-cycle stages (Figure 3), and translate the mechanistic understanding of the system into an integrated model (Figure 4).

By simulating the system at 25m resolution with a daily time step, HELF provides new insight into the time-space patterns of disease risk, which will be valuable for decision support. Compared to the Ollerenshaw Index, which considers each month independently from every other, HELF is dynamic. This means that high rainfall may result into high risk of infection only depending on the antecedent moisture conditions of the soil and their effect on the life-cycle progress (Fig. 12a). Moreover, by providing greater temporal resolution, HELF allows capturing the impact of short-term weather events, such as an extremely warm day or intense concentrated rainfall, which are believed to be particularly relevant for the biological system (13–15). In tandem with the fact that HELF can identify hotspots of transmission potential (Figure 12b), this means it may be possible for farmers to control the magnitude of exposure to fluke in the field, for example by altering management practices to avoid livestock grazing in high risk areas during peak metacercarial abundance. Finally, the stages included in HELF make up the part of the life-cycle which is missing in the model of fluke dynamics within the final host developed in (19). Integration of the two would allow a mechanistic description of the whole cycle, providing the opportunity to assess, for example, the impact of vaccines on infection levels.

In addition to aiding in the tools for the management of fasciolosis, HELF could also benefit the study of other diseases. The same model could be useful for rumen fluke, which is on the rise in British and Irish livestock and has a similar life-cycle to liver fluke, sharing the same intermediate host (43). On the other hand, a different hydrological model component could be employed instead of TOPMODEL, depending on the hydro-environmental drivers relevant for the disease under consideration (3). For example, a model based on freshwater would be needed for diseases involving aquatic intermediate hosts, such as freshwater snails in the case of schistosomiasis (2).

Several assumptions are currently embedded in HELF. Notably, to account for seasonality and spatial aspect of the disease, we assumed that development of the parasite life-cycle is entirely driven by environmental conditions, simplifying the mechanisms related to the intra-molluscan stage and neglecting density-dependent processes. Moreover, even with regard to environmental factors, characteristics such as soil pH and texture have been described as potentially relevant for the presence of snail habitats (27), but have not been included in our model, yet. However, HELF could be expanded to incorporate these, as well as additional spatial data, including remote sensing information.

To address common disease data limitations, we proposed an approach which includes the use of expert knowledge to constrain and evaluate our new model. Fitting observations is standard practice for calibration of hydrologic models, when there is a gauging station providing data to compare simulations against (Figure 6). Distributed soil moisture observations were not available for our case studies (and are rarely available anywhere, especially at high resolution), but previous studies have shown that TOPMODEL can provide good representation of the spatial pattern of saturated areas (44). Less frequently, when data is available, the same is done to parameterise epidemiological models (e.g. 16,45) (10). Our results show that HELF is flexible enough to replicate the observed time-space patterns of infection over the two case study catchments (Figures 7–8). We speculate mismatches remaining when we fit the two datasets are not necessarily due to aspects not yet included in the model only, but may also be related to data issues. The absence of reported cases for 2001 from the Tawe Catchment is believed to have been influenced by the outbreak of foot-and-mouth, which killed over 10 million cows and sheep, affecting submissions to the vet labs. Similarly, discrepancies over sub-areas A2 and A5 in the Severn Catchment may also be related to our underlying assumption of uniform distribution of farms per sub-area, which may not necessarily reflect the real-world system. Mis-reporting and data low space-time resolution are common with many diseases and have often been recognised as a bottleneck to developing models providing meaningful predictions of disease risk (12,14,16). Moreover, even if available epidemiological data were more reliable, they would still reflect historical conditions, which may not necessarily be relevant for the future (11,15). Our calibration strategy includes the use of expert-driven rules to overcome these issues. The rules represent mechanistic knowledge of the system translated into prior information about the output state variables. By using these, we can constrain aspects of the model for which no hard data is available (i.e. the different life-cycle stages) in a process-based manner, without biasing the parameters towards external drivers not included in the model. The current formulation reflects changes in seasonality experienced over our simulation period. However, going forwards, this can be adjusted to account for further changes, in order to reliably assess the impact on disease risk of conditions beyond the range of previously observed variability. Our results show there are parameter sets satisfying all four our rules (Figure 9), and that the resulting behaviour of the simulated stages and lags between them (Fig. 10) agree with what has been traditionally observed in the UK (20,24). This suggests that HELF reflects well (our perception of) the real-world system. The fact that a significant number of simulations is rejected from the initial sample suggests that our parameter confinement strategy is effective, which is crucial as the inability to identify behavioural parameterisations may result in significant predictive uncertainty when using the model under changing conditions (15,34). Moreover, using HELF with Monte Carlo sampling allows explicit consideration of uncertainty, by propagating it from the parameter ranges to the model simulations. This means we can provide decision-makers with a degree of confidence to be attributed to the model results. The reason why uncertainty in the simulated risk of infection still seems high in terms of magnitude (Figure 11) is that the rules are currently based on information about the seasonality of the disease only, driven by our aim of providing a model that is generally applicable in the UK. However, if reliable local data were available, the rules could be modified or increased in number to make the model more accurate locally (as in 16,46). On the other hand, the fact that uncertainty bounds are narrow in terms of timing and duration of the outbreaks is particularly useful to inform farmers’ decisions about e.g. when to allow grazing of animals or when to treat them.

## 7. Conclusions

We developed and tested a new mechanistic hydro-epidemiological model to simulate the risk of liver fluke infection, linked to key weather-water-environmental processes (HELF). The fact that, unlike previous models, HELF explicitly describes the processes, rather than relying on correlation, makes it more likely robust for capturing the impact of ‘new’ conditions on disease risk. We showed that the model is sufficiently flexible to fit observations over two UK case studies, but also introduced an expert-driven calibration strategy to make the model more robust to data with limited reliability and in the presence of climate change. Finally, comparison with a widely-used empirical model of liver fluke risk showed that, while matching the existing index on interannual variation, HELF provides better insight into the time-space patterns of disease, which will be valuable for decision support. Driving the model with climate and management scenarios will enable assessing future risk of infection and evaluating control options to reduce and/or mitigate disease burden. This is urgent, given the widespread increasing drug resistance and threat of altered patterns of transmission due to climate-environmental change.

Through the example of fasciolosis, we demonstrated *(i)* that sufficient mechanistic understanding of the bio-physical system may be available to develop and test a process-based model for an environmentally-mediated disease, without having to rely only on limited and potentially disinformative data, and *(ii)* how accounting for the critical hydro-environmental controls underlying transmission can be valuable to better understand seasonality and spread of emerging or re-emerging threatening diseases.

## Data accessibility

Underlying datasets are publically available and referenced within the paper. The FEC-based dataset is available at http://dx.doi.org/10.17638/datacat.liverpool.ac.uk/406. Underlying code is available at https://github.com/ludobeltrame/helf.

## Authors’ contributions

LB and TW designed research; LB, TD, HRV, ERM, JGW, PW and TW performed research; CMM and DJLW collected the FEC dataset; LB analysed data and wrote the paper with contributions by all other authors.

## Acknowledgements

We thank Sian Mitchell for advising on the VIDA dataset, as well as Philip Skuce, David Armstrong and Nerys Wright, for the input and useful discussions which allowed development of the rules.

